# Age-related Cellular and Microstructural Changes in the Rotator Cuff Enthesis

**DOI:** 10.1101/2021.07.22.452068

**Authors:** Zeling Long, Koichi Nakagawa, Zhanwen Wang, Peter C. Amadio, Chunfeng Zhao, Anne Gingery

## Abstract

Rotator cuff injuries increase with age. The enthesis is the most frequent site of rotator cuff injury and degeneration. Understanding age-related changes of the enthesis are essential to determine the mechanism of rotator cuff injuries, degeneration, and to guide mechanistically driven therapies. In this study, we explored age-related cellular changes of the rotator cuff enthesis in young, mature, and aged rats. Here we found that the aged enthesis is typified by an increased mineralized zone and decreased non-mineralized zone. Proliferation, migration, and colony forming potential of rotator cuff derived cells (RCECs) was attenuated with aging. The tenogenic and chondrogenic potential were significantly reduced, while the osteogenic potential increased in aged RCECs. The adipogenic potential increased in RCECs with age. This study explores the cellular differences found between young, mature, and aged rotator cuff enthesis cells and provides a basis for further delineation of mechanisms and potential therapeutics for rotator cuff injuries.

## INTRODUCTION

The incidence of musculoskeletal system diseases increases with age, resulting in severe joint pain, dysfunction and a large socioeconomic burden^1^. With age musculoskeletal tissues are prone to injury, degeneration, and slow healing.

Shoulder rotator cuff tears are one of the most common musculoskeletal clinical disorders, and the incidence increases rapidly after age 50^1^. There are 4.5 million shoulder injuries per year in the United States, of which 70% are due to rotator cuff tears ^2^. Rotator cuff tears result in chronic pain, tissue dysfunction and reductions in mobility and activity, and thus impair healthy aging^2; 3^.

Rotator cuff injuries are often found at the tendon-bone interface (i.e., the enthesis). The enthesis is a composite tissue with a gradient of tendon, unmineralized fibrocartilage, mineralized fibrocartilage and bone ^1; 4; 5^. This transitional zone from tendon to bone is typified by well-organized tendon fibers splayed out into thinner enthesis fibers which are firmly attached to the bone ^6–10^. This structure exhibits unique functions in flexibility and reducing stress concentration between tendon (soft) and bone (hard) tissue, thus protecting the tendon from fatigue injury at its insertion during shoulder movement^11^.

It is well understood that tendon and bone undergo degenerative processes with age. However, it remains unclear how microstructure and cellular functional properties of the enthesis changes with age. Rotator cuff tears can occur at any age but are increasingly evident with aging ^1; 12^. The repair and restoration of functionality of the enthesis after injury and degeneration remains a significant clinical challenge.

In this study we hypothesized that the cells from rotator cuffs of aged individuals would have reduced proliferation, migration, tenogenic and osteogenic potential, as compared to cells from younger individuals. To explore this, we used a rat shoulder rotator cuff model. We isolated rotator cuff enthesis cells (RCECs) from young, mature, and aged rats. RCECs are a mixed population of cells that are found in the rotator cuff including tenogenic, chondrogenic and osteogenic cells. To our knowledge this is the first time that primary cells isolated from young, mature, and aged rotator cuff entheses have been assessed for proliferation, migration, colony formation and progenitor potential.

## METHODS

### Sample preparation

Supraspinatus-enthesis-bone complex (supraspinatus were used here for its particular vulnerability in aging process ^13^) were dissected from male Sprague Dawley rats immediately after sacrifice from other Institutional Animal Care and Uses Committee-approved studies. All samples were divided into three groups according to the rats’ age: Young adult group (Y group 8-12 weeks), Mature group (M group 12-15 months) and Aged group (A group 20-24 months).

### Histology and image analysis

Five samples from each group were fixed in 4% paraformaldehyde and decalcified in decal solution for 10 days, then embedded in paraffin and cut into ~7μm thick slide. H-E, Toluidine Blue and Picrosirius red staining were conducted following manufacturer’s protocols. Slides were observed with 40 times magnification under a light microscope and images were collected by a camera mounted on the microscope. All measurements were performed by two independent researchers. One H&E stained slide and one Toluidine blue stained slide from each sample were used for cell counting and area measurement. Briefly, the enthesis was divided into mineralized and non-mineralized zone by the tidemark, the region between tidemark and subchondral bone is mineralized zone and the region distal to the tidemark is non-mineralized zone. Both regions were outlined according to these criteria. Numbers of cells in each region were manually counted using Photoshop counting tool. Areas of mineralized/ non-mineralized region were quantified. For Picrosirius red staining, a polarized light scope was used to observe collagen fibers. Ten square regions were randomly selected from each slide and the grayscale value was measured using Image J software.

### Rotator cuff enthesis derived cells isolation

Isolation of rotator cuff enthesis derived cell (RCECs) based on the method of Harada et al with minor modifications ^14; 15^. Briefly RCECs were derived from four different individuals in each group. Immediately after sacrifice, the supraspinatus and infraspinatus tendon were cut off along the tendon-bone conjunction. The conjunction tissue was cut into small fragments and cultured in 75 mm plate with Dulbecco’s modified eagle medium (DMEM, Thermo Fisher) containing 10% fetal bovine serum (FBS, Sigma) and 1% antibiotics at 37 C with 5% humidified CO2. After two weeks of culture, the adherent cells attached, the fragments were removed, the culture medium was changed every three days and passaged when they reach 80% confluent until passenger three (P3).

### Proliferation and migration assay

The proliferation and migration assays were conducted to evaluated. Briefly, RCECs (n=4) were triple-plated in12 well plates at 5×10^4^/well and incubated in IncuCyte (IncuCyte S3 Live-cell analysis system) with automated image acquisition every 8 hours until confluence. A scratch was made using 100ul tips on each well and the migration and healing process was recorded by the IncuCyte every 8 hours for two days. Four locations were randomly selected to measure the healing area. The culture medium was changed every three days.

Cell counting kit-8 (Dojindo) assay was used to verify the proliferation assay result of IncuCyte. Briefly, RCECs were seeded in 96-well plates at a concentration of 1000 cells per well. Cells were stained with CCK-8 kit following the manufacturer’s protocol. OD value at 450nm at day one, three and five were collected.

### Colony-formation Unit (CFU) assay

The colony-forming unit assay was conducted to evaluate self-duplicate capacity. Briefly, RCECs (n=4) were triple plated on 60mm plates in triplicate at 100 cells per dish with media half-changed every three days. After ten days of cultivation, RCECs were fixed with 4% PFA for 15 minutes and stained with crystal violet for 10 minutes (Sigma, St. Louis, MO) and counted using a light microscope.

### Osteogenic differentiation

RCECs (n=4) were triple-plated in 12-well plates at a concentration of 5×10^4^ cells/well and cultured with osteogenic differentiation medium (Gibco; StemPro ® Osteogenesis Differentiation Kit). The cells cultured in basal culture medium served as controls. Culture medium was changed every three days. After 21 days of culture, osteogenic differentiation was evaluated by Alizarin red S staining of the mineralized matrix, lineage-specific genes (Osteopontin (Spp1), RUNX Family Transcription Factor 2 (Runx2), Osteocalcin (Bglap), primers sequences are listed in Supplementary 1) were evaluated by q-RT PCR.

### Adipogenic differentiation

RCECs (n=4) were triple-plated in 12-well plates at a concentration of 5×10^4^ cells/well and cultured with adipogenic differentiation medium (Gibco; StemPro ® Adipogenesis Differentiation Kit). The cells cultured in basal culture medium served as controls. Culture medium was changed every three days. After 21-days-culturing, adipogenic differentiation was evaluated by Oil red-O staining of the cellular accumulation of neutral lipid vacuoles. Lineage-specific genes (Peroxisome Proliferator Activated Receptor Gamma (Pparg), adipocyte Protein 2 (Ap2), Lpl (lipid transfer protein 1), primers sequences are listed in Supplementary 1) were evaluated by q-RT PCR.

### Chondrogenic differentiation

An aliquot of 8×10^5^ RCECs (n=4, 3 repeats) was spun down at 1500rmp for 3 min in a 15-ml polypropylene tube to form a pellet. Cell pellets were cultured with chondrogenic differentiation medium (Gibco; StemPro ® Chondrogenesis Differentiation Kit). The cells cultured in basal culture medium served as controls. Culture medium was changed every three days. After 21 days of culturing, chondrogenic differentiation was evaluated by toluidine blue staining, the pellets were embedded in OCT and cryo-sectioned for microscopy. Lineage-specific genes (collagen type II (Col-IIa1), SRY-Box Transcription Factor 9 (Sox9), collagen X (Col-X), primers sequences are listed in Supplementary 1) were analyzed by q-RT PCR.

### Tenogenic differentiation

RCECs (n=4) were triple-plated in 12-well plates at a concentration of 5×10^4^ cells/well. and cultured with a supplement of 50ng/ml BMP12 (catalog# 120-37, peproTech). The cells cultured in basal culture medium served as controls. Culture medium was changed every three days. After 14-days-culturing, tenogenic differentiation was evaluated by Picrosirius red staining of collagen deposition. Lineage-specific genes (Tenomodulin (Tnmd), Tenascin (Tnc), Decorin (Dcn), primers sequences are listed in Supplementary 1) were evaluated by q-RT PCR.

### Gene Expression

For gene expression analysis total RNA was isolated by Trizol (Invitrogen Life Technologies; Thermo Fisher Scientific), and the RNA was reverse transcribed into cDNA using an iScriptTM cDNA Synthesis Kit (Bio-Rad Laboratories, Hercules, CA, USA). Quantitative PCR was performed with RT2 SYBR Green qPCR Master Mix (QIAGEN, Hilden, Germany), using an AC1000 Touch™ Thermal Cycler (Bio-Rad Laboratories) under the following cycling conditions: 95 °C for 10 min followed by 40 cycles of amplification (95 °C for 15 s and 60 °C for 1 min). Expression levels were geomean normalized to β-actin, GAPDH, and TBP. Samples were compared using ΔΔCt method.

### Statistical Analysis

Numerical data are presented as mean ± standard deviation and were tested for normality. Statistical analyses of the histological quantification were performed using Kruskal-Wallis test. Two-way analysis of variance (ANOVA) and multiple comparisons were used to compare the CCK8, proliferation, migration, CFU and gene expression results among groups. Relative gene expressions were normalized to young non-differentiated group. Multiple comparison was conducted in two ways, 1) between young non-differentiated group and every other groups, 2) between each pair of differentiated group and non-differentiated group. All statistical analyses were performed with SPSS software (version 8.2.1; SPSS Inc); a P value of <0.05 was considered significant.

## RESULT

### Histological Age Changes in Rotator Cuff

Histological assessment of enthesis morphology, cellular fraction, mineralization, and collagen alignment were assessed with H&E, Toluidine blue and Picro Sirius staining. Samples were divided into three groups according to age (young group: 8-12 weeks, mature group: 12-15 months, aged group: 20-24 months). In the young group the margins of the enthesis are well defined with small and well-aligned cells. The tide mark dividing mineralized area and non-mineralized area is clear and sharp (Figure 1A), large defined polysaccharide regions were found in toluidine blue stained sections (Figure 1D). In the mature group, the structure of enthesis remained intact, but with increased cellular disorder and decreased polysaccharide expression (Figure 1B&E). In the aged rotator cuff the cellular organization and tidemark were increasingly disorganized and the fibro-chondrocytes in mineralized area enlarged and disorganized. The polysaccharide region is substantially reduced (Figure 1C&F). Quantification of histological data is shown (Figure 1J). Data are reported as mean ± SD. Cell counts in the mineralized area are 372.7 ± 202.87, 220.4 ± 63.11 and 383.3 ± 159.06 in young, mature, and aged groups respectively (P=0.0362). In the non-mineralized area, the cell counts are 401 ±197.14, 260 ±100.88 and 182.5 ± 89.23 young, mature, and aged groups respectively (P=0.0094). Mineralized areas in young, mature, and aged groups are 0.069 ± 0.024 mm^2^, 0.042 ± 0.015 mm^2^ and 0.072 ± 0.023mm^2^ (P=0.0037) while non-mineralized areas are 0.072 ± 0.020 mm^2^, 0.059 ± 0.017 mm^2^ and 0.031 ± 0.007 mm^2^ respectively (P=0.0001) (Figure 1K). The thickness of the enthesis was measured as the mean of three randomly selected locations. Mineralized thickness in these three groups are 0.092 ± 0.026 mm, 0.055 ±0.011 mm and 0.125 ±0.027 mm (P<0.0001) in the young, mature, and aged groups. Non-mineralized area, they are 0.109 ±0.021 mm, 0.083 ±0.021 mm and 0.057 ±0.019 mm (P=0.0004) respectively (Figure 1L). Brightness greyscale values were measured using Image J software based of picrosirius red staining figures (Figure 1GHI). They are 61.51 ± 23.54, 63.41 ± 25.14 and 69.38 ± 31.71 for young, mature, and aged groups (P=0.1554).

**Figure 1.**
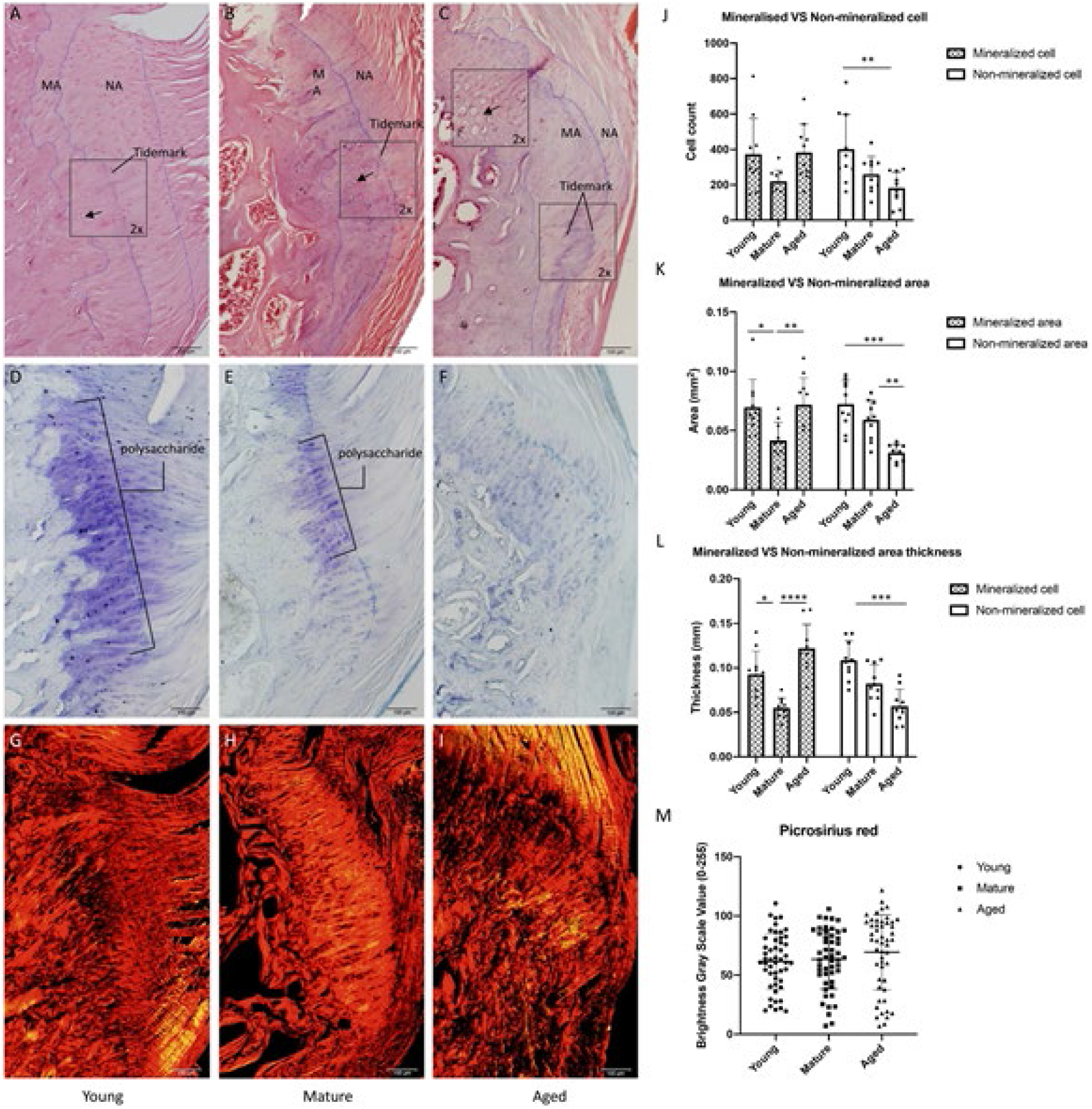
Histology of rotator cuff enthesis in young, mature and aged rats. Rotator cuff enthesis was stained with Hematoxylin & Eosin (H&E) for cellular structure (A, B, C). Mineralized area (MA) and non-mineralized area(NA) were divided by tidemark. The mineralized cells are small and distributed in pairs in young group but become larger and clustered with aging (black arrows), tidemark becomes rougher and deeper stained with age; Toluidine blue (TB) staining indicates decreasing polysaccharides with age (D, E, F). Cell counts and measures of areas were quantified based on H&E staining and TB staining. Non-mineralized area (K, L) and cells (J) decrease with age. Mineralized area and cell number show the lowest value in mature group. Brightness gray scale value (M) based on Picrosirius red staining (G, H, I) indicates collagen alignments decrease with age, though it did not reach significance. (Young: 8-12 weeks, Mature: 12-15 months, Aged: 20-24 months). * P <0.05, ** P<0.01, *** P <0.001, **** P<0.0001.

### Proliferation and wound healing assay

Rotator cuff enthesis cells (RCECs) were isolated and cultured from young, mature, and aged rats. Cells were cultured and assessed using proliferation and migration assays. After 10-14 days of culture, radial clustered, spindle-shaped cells were observed, exhibiting a fibroblast-like morphology. This feature is clear in young group (Figure 3A) but diminished with age. In the aged group, the RCECs are rounded, enlarged and isolated morphology (Figure 3 C).

Next, we assessed proliferation potential of passage three RCECs. Here we found that young RCECs showed a significantly higher proliferation rate as compared to mature and aged RCECs as evaluated by CCK8 proliferation assay (Figure 2B) and fold change assessed by IncuCyte phase object confluence analysis (Figure 2C).

**Figure 2.**
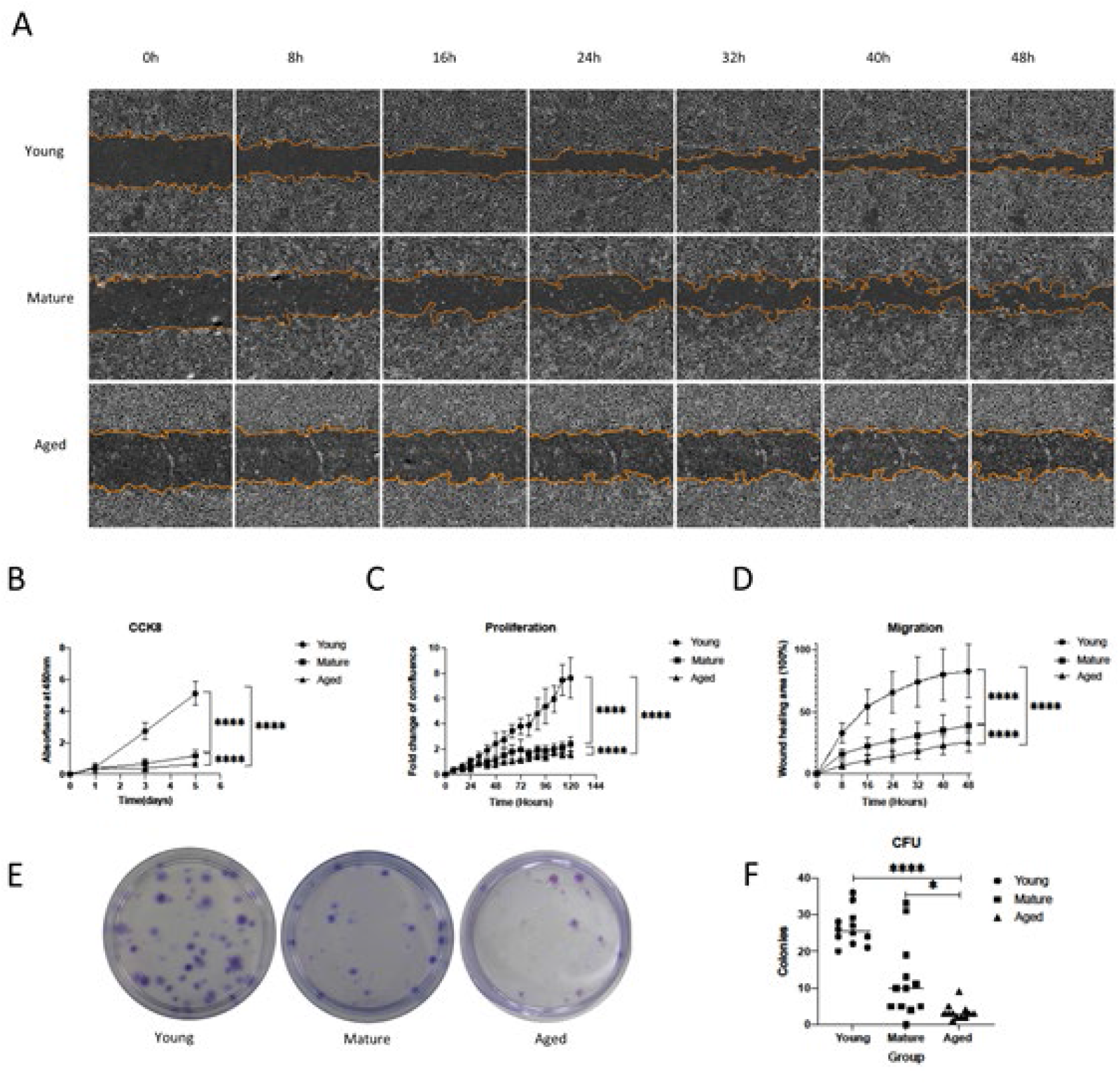
Proliferation and Migration assays of RCECs. Proliferation assays were performed with CCK8 proliferation assay method (B) and IncuCyte recording the fold change of confluence (C), significance was shown in both assays between all the age groups. Migration (D) assay results were illustrated as fraction of wound healing area. Young RCECs demonstrated significantly higher migration ability than mature and aged RCECs. Colony formation Units (E) were stained with crystal violet staining for visualization, more units were observed in young group (F) and these units are larger than that of mature and aged group. (Young: 8-12 weeks, Mature: 12-15 months, Aged: 20-24 months) * P <0.05, **** P <0.0001.

We further assessed RCECs migration potential. Our migration assay shows that young RCECs have significantly increased migration rate as compared to mature and aged cells. After 48 hours of culture, seven out of sixteen selected areas fully bridged the scratched region in young RCECs cultures, compared to the mature RCEC’s highest migration (of 59% healing area) and aged RCECs of 40% healing area (Figure 2AD).

### Colony-forming Units

Colony forming units (CFU) are another measure of proliferative potential. The proportion of cells with self-replicating potential was determined by counting the number of colonies, to evaluate the overall cell proliferation potential. Here we assessed the ability of RCECs to form CFU (Figure 2E). Young RCECs formed significantly more CFUs than mature and aged RCECs. The mean colony formation unit for young RCECs was 26.33 ± 4.89 units per 100 cells, significantly higher than both the mature (12.17 ± 10.51 units per 100 cells, P<0.01) and aged groups (3.33 ± 2.06 units per 100 cells, P<0.001) (Figure 2F). Further we noted that the size of colonies formed in young group was larger than those in mature and aged RCECs.

### Multi-differentiation Potential

RCECs in all three age groups had the ability to differentiated in osteogenic, adipogenic and chondrogenic lineages. However, mature RCECs and aged RCECs did not differentiation to tenogenic lineage in contrast to young RCECs.

Calcium accumulation, demonstrated by Alizarin red staining, was formed in all ages of osteogenic RCECs (Figure 3DEF). Lineage-specific osteogenic genes (Spp1, Runx2, Bglap) were up-regulated in all differentiated groups, interactions between age and treatment (osteogenesis) were significant (Figure 3GHI).

**Figure 3.**
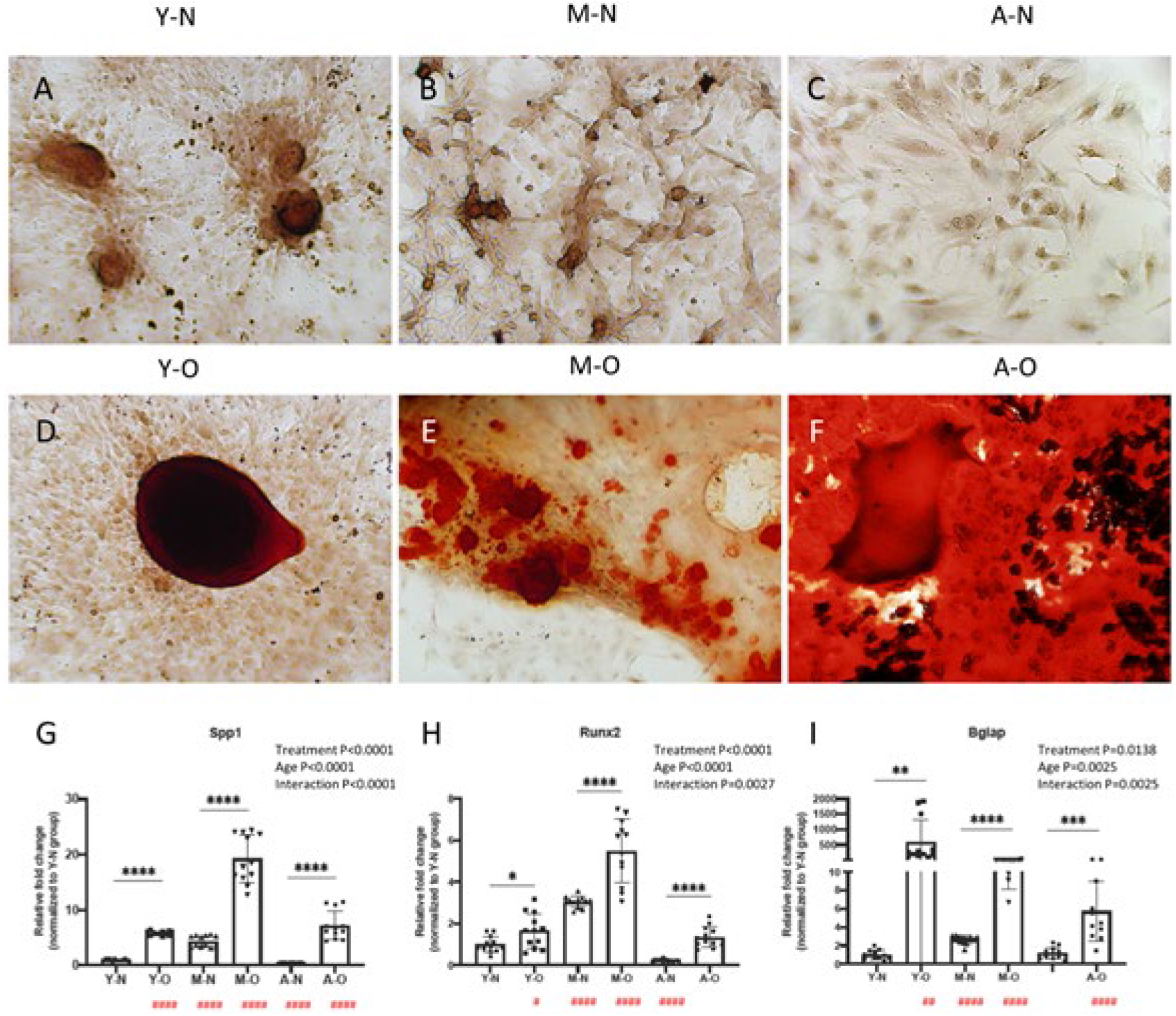
Osteogenic potential of RCECs at different ages. RCECs from different-age groups were successfully differentiated into osteocytes lineage under inducement. Red-stained calcium accumulation was formed in differentiated RCECs (DEF). The expression of osteogenic lineage specific gene was significantly increased (G-I). Treatment: the independent effect of inducement on gene expression. Age: the independent effect of age on gene expression. Interaction: the interaction of inducement and age on gene expression. Y-N: Young-nondifferentiated; Y-O: Young-osteogenesis; M-N: Mature-nondifferentiated; M-O: Mature-osteogenesis; A-N: Aged-nondifferentiated; A-O: Aged-osteogenesis; */# P<0.05, **/## P<0.01, ***/###P<0.001, ****/#### P<0.0001, # represents the significant differences with Y-N group.

Adipogenesis was verified by Oil red staining, red-stained lipids were found in all ages of adipogenic RCECs (Figure 4DEF). Lineage-specific adipogenic genes (Pparg, Ap2, Lpl) were up-regulated in all differentiated groups, interactions between age and treatment (adipogenesis) were significant (Figure 4 GHI).

**Figure 4.**
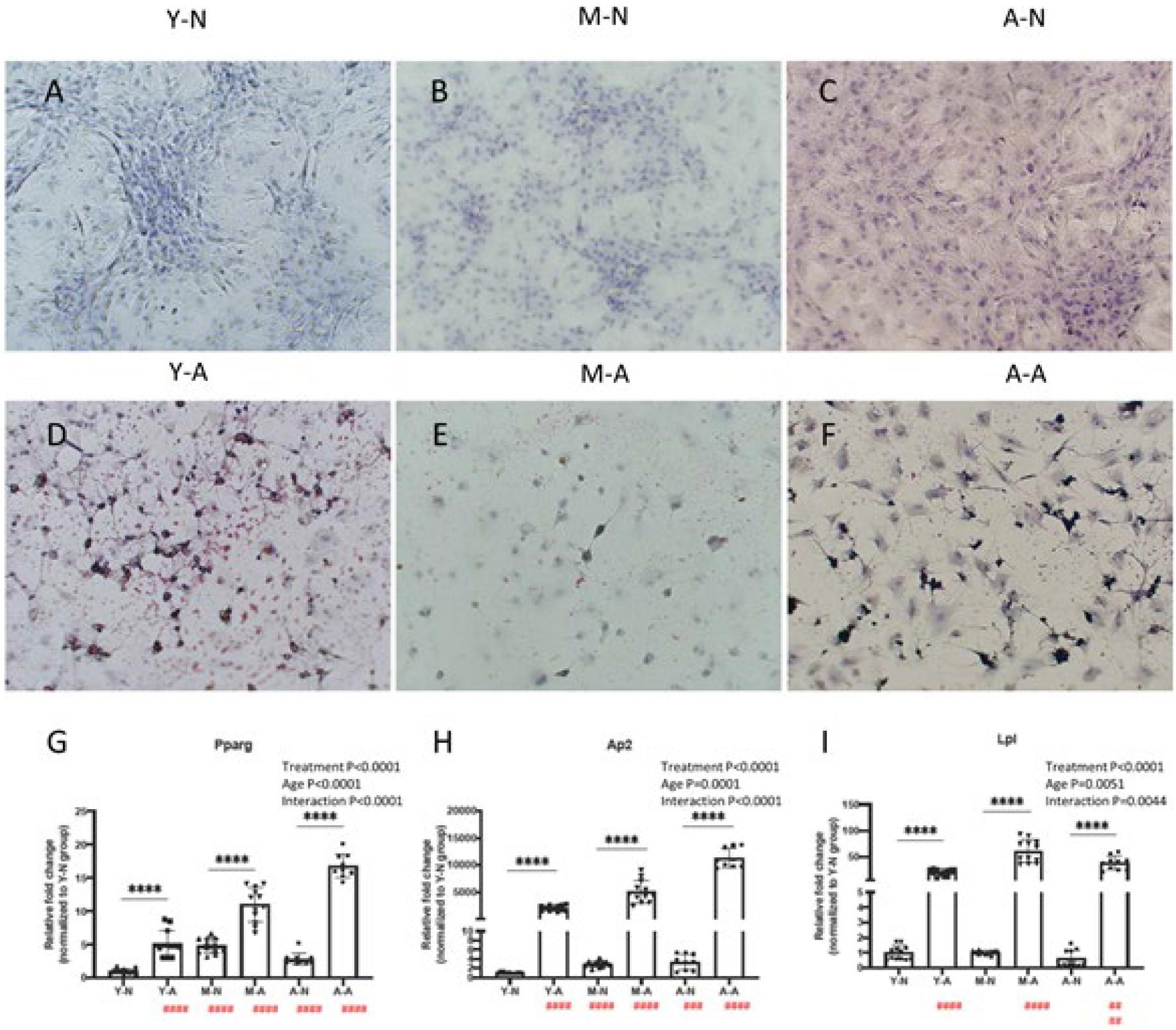
Adipogenic potential of RCECs at different ages. RCECs from different-age groups were successfully differentiated into adipocytes lineage under inducement. Red-stained lipids were formed in differentiated RCECs (DEF). The expression of adipogenic lineage specific gene was significantly increased (G-I). Treatment: the independent effect of inducement on gene expression. Age: the independent effect of age on gene expression. Interaction: the interaction of inducement and age on gene expression. Y-N: Young-nondifferentiated; Y-A: Young-adipogenesis; M-N: Mature-nondifferentiated; M-A: Mature-adipogenesis; A-N: Aged-nondifferentiated; A-A: Aged-adipogenesis; */# P<0.05, **/## P<0.01, ***/### P<0.001, ****/#### P<0.0001, # represents the significant differences with Y-N group.

Chondrogenesis of RCECs was assessed by Alcian blue staining and mRNA expression of lineage-specific genes (Collagen II, Collagen X, Sox9). Before chondrogenic inducement, RCECs demonstrated a connected, reticular fibroblast-like morphology. After 21 days of chondrogenic treatment, RCECs exhibited rounded, granular chondrocyte-like phenotype in all ages of chondrogenic groups (Figure 5DEF). Lineage-specific chondrogenic genes were up-regulated in young and mature chondrogenic RCECs. However, Col-II and Sox9 were not up-regulated in aged chondrogenic RCECs, only COL-X was up-regulated in aged chondrogenic RCECs. Interactions between age and chondrogenic treatment were significant (Figure 5GHI).

**Figure 5.**
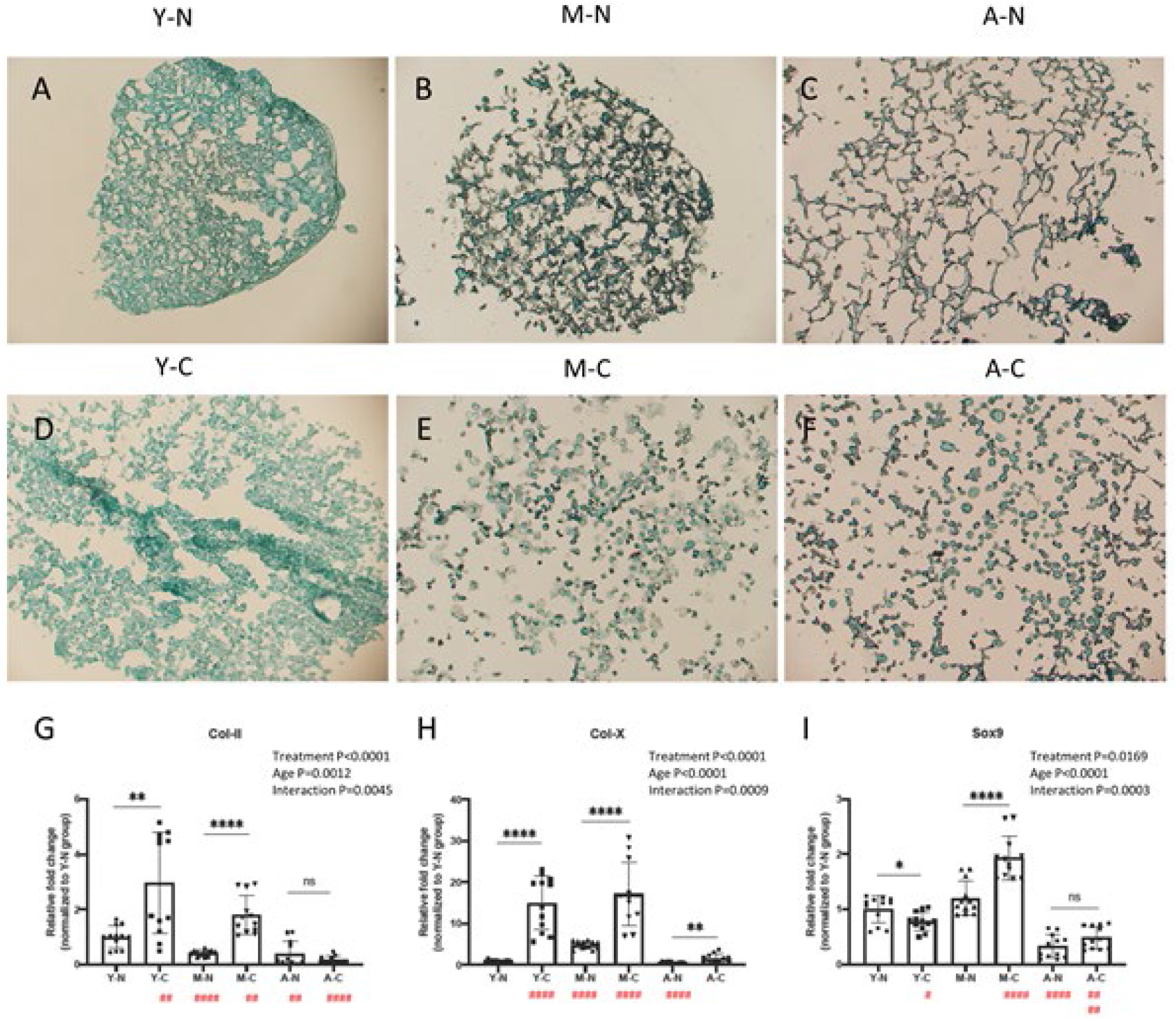
Chondrogenic potential of RCECs at different ages. RCECs from different-age groups were successfully differentiated into chondrocytes lineage under inducement. RCECs exhibited rounded, granular chondrocyte-like phenotype (DEF). The expression of chondrogenic lineage specific gene was significantly increased (G-I). Treatment: the independent effect of inducement on gene expression. Age: the independent effect of age on gene expression. Interaction: the interaction of inducement and age on gene expression. Y-N: Young-nondifferentiated; Y-C: Young-chondrogenesis; M-N: Mature-nondifferentiated; M-C: Mature chondrogenesis; A-N: Aged-nondifferentiated; A-C: Aged-chondrogenesis; */# P<0.05, **/## P<0.01, ***/### P<0.001, ****/#### P<0.0001, # represents the significant differences with Y-N group.

RCECs were treated with BMP12 for 14 days a noted tenogenic inducer ^16–18^. Picrosirius red collagen staining was performed to visualize the tenogenic differentiation. Young RCECs had strong red collagen staining, and which was enhanced after BMP12 induction (Figure A, D). However, there was less staining in the mature and aged groups, nor after BMP12 treatment. The aged RCECs displays enlarged, rounded, isolated morphology before treatment (Figure 6C), but become more spindle-shaped and clustered after BMP12 induction (Figure 6F). Lineage-specific gene Tnmd expressed significantly higher in the young RCECs with or without linage induction (Figure 6G) and were up-regulated in BMP12 tenogenic RCECs induction in the mature and aged groups. Interestingly, for Tnmd, there were no significant interactions between age and treatment, while both separately were significant. Tnc, interactions between age and treatment were not significant (Figure 6H). The expression of Dcn was up regulated after BMP12 induction in young RCECs and aged RCECs. Expression of Dcn was much lower in mature and aged RCECs regardless of BMP12 induction. Interactions between age and treatment were significant (Figure 6I).

**Figure 6.**
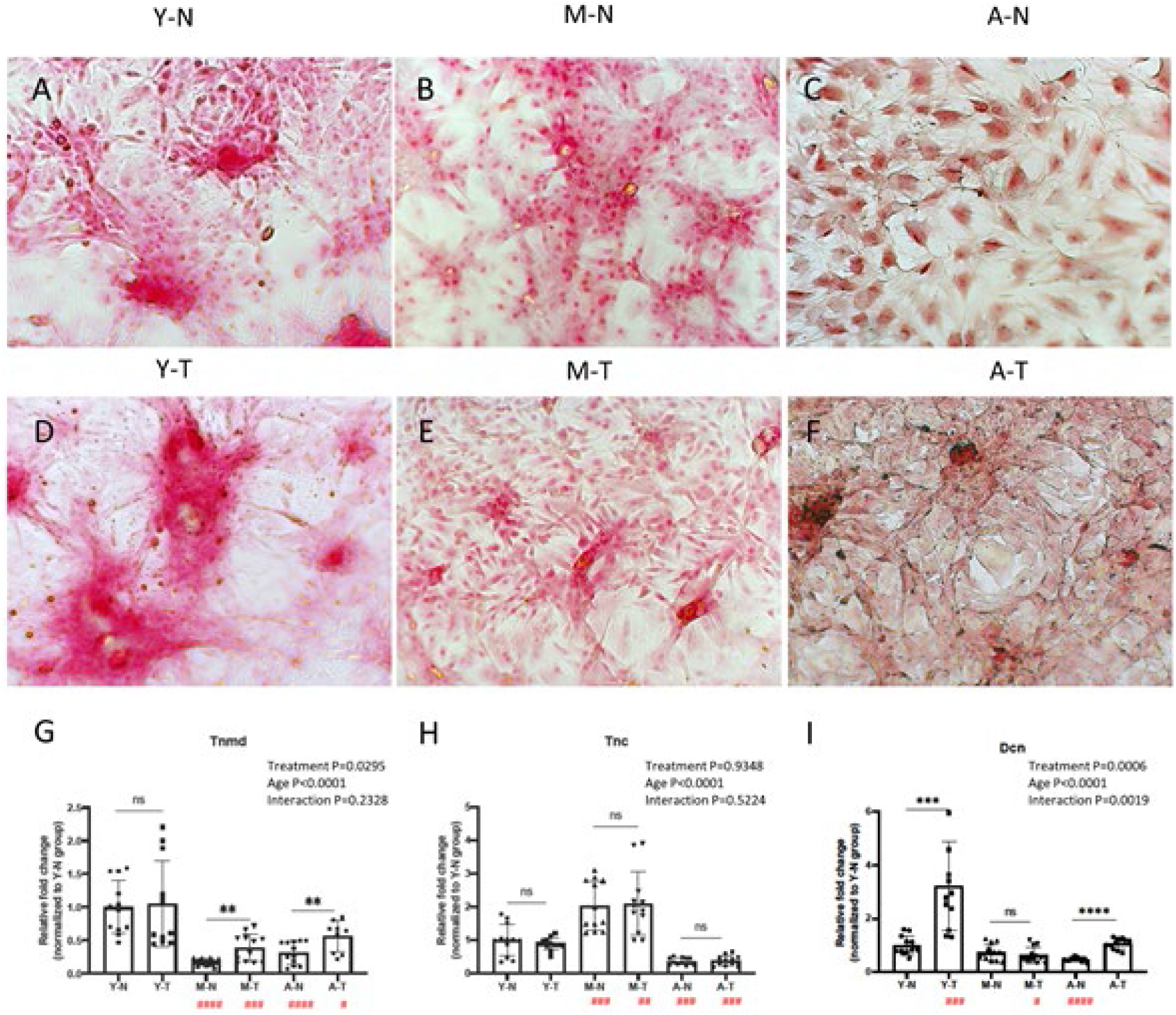
Tenogenic potential of RCECs in different ages. RCECs from different-age groups were successfully differentiated into tenocytes lineage under inducement. Young RCECs show strong collagen staining before and after tenogenic induction (A&D). However, there was less red stained collagen matrix in the mature and aged groups (BC&EF). The expression of chondrogenic lineage specific gene Tnmd was significantly higher in young groups, tenogenic inducement up regulated the expression of Tnmd in all groups, though there is no significance in young group (G). Dcn was upregulated significantly in young differentiated group and aged group. Treatment: the independent effect of inducement on gene expression. Age: the independent effect of age on gene expression. Interaction: the interaction of inducement and age on gene expression. Y-N: Young-nondifferentiated; Y-T: Young-tenogenesis; M-N: Mature-nondifferentiated; M-T: Mature-tenogenesis; A-N: Aged-nondifferentiated; A-T: Aged-tenogenesis; */# P<0.05, **/## P<0.01, ***/### P<0.001, ****/#### P<0.0001, # represents the significant differences with Y-N group.

## DISCUSSION

This study explored the age-related cellular and microstructural changes found in the rotator cuff enthesis using a rat model. Histological assessment showed the typical gradient structure of entheses in our models including the fibrous structure of tendon, non-mineralized fibrocartilage, and mineralized fibrocartilage and bone. With age well-aligned collagen fibers in the rotator cuff enthesis become increasingly disorganized. Further with age, this enthesis zone narrows, substantially decreasing the contact area between tendon and bone.

There is a clear boundary between bone tissue and fibrocartilage in the enthesis, which is more obvious with toluidine blue staining. In fibrocartilage, the polysaccharide in the matrix stains blue, which is not seen in the bone area. Chondrocytes/fibro-chondrocytes synthesize polysaccharide. Polysaccharides allow the ECM to be permeable and able to absorb more water, which helps to resist compressive loads. With age the polysaccharide area diminishes, resulting in poor mechanical performance.

The fibrocartilage is further divided into mineralized and non-mineralized areas, separated by the tidemark^19^. Cells in mineralized fibrocartilage are more differentiated, rounded and hypertrophic^20; 21^, while in non-mineralized fibrocartilage, they are smaller and aligned in columns, similar to immature chondrocytes^22; 23^. In our findings, the non-mineralized fibrocartilage and cellularity decrease with age. The non-mineralized fibrocartilage plays an important role in dissipating the stress upon loading ^24–26^, so a decrease in the size of this zone may result in a greater risk of injury. Further the reduced number of cells in non-mineralized fibrocartilage may indicate reduced regeneration potential. Besides, the tide mark becomes increasing disorganized with age, which reflect the resultant tears and attenuated mechanical properties ^27^. Collectively, these structural changes weaken the elastic buffer effect of the enthesis, making it more fragile and prone to injury.

Structural changes are mitigated in large part by cellular changes that contribute to the rotator cuff injuries that are found with aging. There are three dominating aging cellular changes, a) decreased proliferation rate, b) loss of wound-healing/migration capability and c) shifted differentiation potential.

Uniquely in this study, we isolated primary cells from the rotator cuff enthesis, instead of focusing on specific cell types, as this more accurately represents the cellular diversity found in the enthesis. The cell types can include cells from bone, cartilage, fibrocartilage, tendon, stromal progenitors and adipose tissue. Assessing the different lineages separately limits the ability to assess the overall impact of this composite tissue. The assessment of the primary enthesis cells provides a broader picture of enthesis as an integrated tissue, which is the functional unit of the tendon-bone interface. The primary RCECs were assessed for migration, proliferation, and differentiation potential. Here we found that, RCECs derived from mature and aged rats have significantly decreased proliferative, colony forming, and migrating potential as compared to the RCECs from young animals. In the young enthesis when micro-wounding occurs^28^, the RCECs can quickly proliferate and repair the damage, so as to avoid further tears of the rotator cuff. However, in the aged enthesis, this ability is weakened, the damage cannot be repaired effectively in time, consequently the rotator cuff tear can more readily propagate.

Assessment of RCECs in different age groups showed that each group retained their multi-lineage differentiation potentials, however the potential for differentiation shifted. Significant interactions were found between treatment and age in these differentiation experiments. Mature and aged RCECs demonstrated high differentiative potential towards osteocyte and adipocyte, as shown in the lineage specific gene expression and histological staining.

After chondrogenic differentiation, lineage specific markers of RCECs increased significantly in young and middle-aged groups, but not in aged group. Only Col-X was up-regulated in aged chondrogenic RCECs. This type of collagen is synthesized by hypertrophic chondrocytes, which is evident in aged chondrocytes^20; 21; 29^.

Assessing the role of tenogenic lineage capacity is more challenging. To date there is no established method for this tenogenic assessment. Some reports have suggested tenogenic induction using BMP12 ^16–18^. Although mature and aged RCECs had some tenogenic potential, tenogenic markers were largely attenuated in mature and aged RCECs. It is important to note that among all tenogenic markers, Tnmd tends to be the most representative biomarker for stemness ^30–33^. The significant loss of Tnmd in mature and aged RCECs is an essential indicator of the loss tenocyte related gene expression and perhaps differentiation ^34; 35^. Further, the loss of Tnmd can cause premature tendon aging by dysregulating the collagen fibrinogenesis and reducing cell proliferation ^36 37^. Interestingly, Tnmd was not up regulated in young RCECs after BMP12 induction. It is possible that RCECs in young group show a strong phenotype of tendon/tendon stem cells already. In previous studies, Tnc was used as a marker for tendon differentiation^38; 39^, but Tnc is not solely expressed in tendon, being highly expressed in bone and cartilage as well ^40; 41^. Our results show that Tnc was not up regulated after BMP12 induction.

The results of the multi-differentiation assays showed that in the aging process, the RCECs gradually lose their chondrogenic and tenogenic potentials. This makes it difficult to form the typical enthesis structure with four distinct zones in the process repair and regeneration, especially the fibrocartilage structure.

The age-related phenotypic changes of mixed cell populations at the enthesis have not been assessed in previous studies, to the best of our knowledge. Understanding cellular changes that occur with age will aid in the development of novel targets and therapeutic approaches to improve tendon to bone healing that is required for clinical repair including shoulder rotator cuff injuries. A possible strategy to attenuate the damage caused by aging would be the shifting of the aged osteogenic/adipogenic enthesis phenotype to a more fibrogenic phenotype. Approaches to change cellular phenotypes include cell fusion^42^, nuclear transfer^43^, and micro vesicles-based therapy ^44^. These cell-derived vesicles have been shown to alter cellular phenotypes in many different cells and tissue combinations ^45–49^. Our future work will focus on determining if promoting fibro-tenogenic differentiation will aid in the regeneration in aged rotator cuff enthesis which may help reduce age-related injuries and improve native tissue regeneration after repair.

During aging the cells of the rotator cuff enthesis showed declined migration, proliferation and colony forming potential. Further with age the cells of the enthesis have increased adipogenic and osteogenic induction, whereas tendon markers are reduced. These results correspond to the increased fatty osteogenic infiltration found in the enthesis with aging. Taken together these cellular shifts within the enthesis may be responsible for both the frequent rotator cuff injuries and the limited healing and regeneration that is found in patients as they age.

## Acknowledgements

This study was funded by a NIH/NIAMS AR07381 grant, Mayo Clinic Orthopedic Surgery Career Development grant, China Scholarship Council, and the Mayo Clinic Foundation.

